# Auxin signaling in the cambium promotes tissue attachment and vascular development during *Arabidopsis thaliana* graft formation

**DOI:** 10.1101/2023.07.02.547393

**Authors:** Phanu T. Serivichyaswat, Abdul Kareem, Ming Feng, Charles W. Melnyk

**Author notes:** Correspondence: Charles W. Melnyk.

## Abstract

The remarkable ability of plants to regenerate wounds is exemplified during the process of plant grafting when two plants are cut and joined together to grow as one. During graft healing, damaged tissues attach, cells proliferate and the vasculatures connect to form a graft union. The plant hormone auxin plays a central role and mutants perturbed in auxin response fail to successfully graft. Here, we investigated the role of individual cell types and their response to auxin during *Arabidopsis thaliana* graft formation. By employing an inducible misexpression system, we blocked auxin response in individual cell types using the *bodenlos* mutation. We found that auxin signaling in procambial tissues was critical for both successful tissue attachment and also for vascular differentiation. In addition, we found that auxin signaling was required for cell divisions of the procambial cells during graft formation. Loss of function mutants in cambial pathways also perturbed attachment and phloem reconnection. We propose that cambium and procambium are key tissues that allow both tissue attachment and vascular differentiation during successful grafting. Our study thus refines our knowledge of graft development and furthers our understanding of regeneration biology and the function of cambium.

## Introduction

Regeneration in response to injuries is vital for survival of organisms. Plants have a robust regenerative ability that rapidly and efficiently heal wounds and reconnect tissues following damage (Ikeuchi et al., 2019). Since ancient times, we have taken advantage of such regenerative abilities to join the shoot segment (scion) of one plant to the root segment (rootstock) of another, a technique called grafting (Mudge et al., 2009). Grafting has allowed us to improve plant traits and yields by combining superior properties such plant size, biotic and abiotic tolerance into one plant (Melnyk, 2017c). Upon cutting and joining of tissues, graft formation occurs through a series of regenerative events including tissue adhesion, formation of an undifferentiated cell mass called callus, and vascular differentiation between scion and rootstock (Jeffree & Yeoman, 1983; Flaishman et al., 2008; Melnyk et al., 2015). Tissue adhesion is facilitated by cell-wall modifying enzymes (Pitaksaringkarn et al., 2014; Notaguchi et al., 2020; Frey et al., 2022) whereas epidermal and cortical cells expansion during graft healing fills the wound (Melnyk et al., 2015; Matsuoka et al., 2016). Cell divisions occur at the region of tissue adhesion, particularly in the vascular tissues, and are important for successful graft formation (Melnyk et al., 2015; Matsuoka et al., 2016). Wound induced callus also forms and is thought to arise from vasculature and pericycle cells (Ikeuchi et al., 2017). Cambium too plays an important role during graft formation. Wounding induces divisions in the cambium and mutations in several genes important for cambium formation, including DOF and ANAC transcription factors, reduce both grafting efficiency and cambium formation (Matsuoka et al., 2021; Zhang et al., 2022). In addition, grafting handbooks highlight the importance of cambial alignment for grafting (Garner & Bradley, 2013). Thus, multiple cell types likely play an important role during grafting.

One such cell type is procambium, and during *Arabidopsis thaliana* development, procambium formed during primary growth undergoes periclinal divisions to form cambium (De Rybel et al., 2016). Several genes and pathways are known to contribute towards procambium and cambium formation (Shi et al., 2019). For instance, *PXY* and *WOX4* define differentiating xylem cells and cambium cells close to the xylem (Shi et al., 2019). *SMXL5* defines differentiating phloem cells and cambium cells close to the phloem (Shi et al., 2019). Such vascular differentiation of the cambium requires auxin signaling and is a hallmark of secondary growth (Smetana et al., 2019; Mäkilä et al., 2023). Auxin is well known for its role in promoting vascular development and patterning (Wetmore & Rier, 1963; Sachs, 1969). Auxin relies on perception by the TRANSPORT INHIBITOR RESPONSE1/ AUXIN SIGNALING F-BOX (TIR1/AFB) receptors which degrade the Auxin/INDOLE-3-ACETIC ACID (Aux/IAA) proteins to allow AUXIN RESPONSE FACTOR (ARF) activation (Chapman & Estelle, 2009). Auxin plays an important role during graft formation and perturbing auxin response or inhibiting auxin transport blocks grafting (Melnyk et al., 2015; Matsuoka et al., 2016). One example is a gain-of-function mutation in *INDOLE-3-ACETIC ACID INDUCIBLE 12* (*IAA12*)/*BODENLOS* (*BDL*) that inhibits phloem reconnection at the graft junction (Melnyk et al., 2015; Serivichyaswat et al., 2022). *bdl* blocks the auxin signaling pathway with a point mutation in its degradation domain, which prevents the binding of TIR1/AFB, and therefore prevents bdl protein degradation and ARFs remain sequestered (Hamann et al., 2002; Figueiredo & Strader, 2022). Investigating how auxin and *bdl* affect grafting is difficult because *bdl* is constitutively expressed in many cell types and the mutation, along with many auxin mutants, has pleiotropic effects, particularly on embryonic root formation (Hamann et al., 2002). Here, we investigated the role of individual cell types during graft formation and their response to auxin. We employed a chemically inducible gene expression systems to blocked auxin signaling in different cell layers by misexpressing *bdl*. We find that auxin signaling in the cambial tissues is required for both tissue attachment and vascular differentiation, whereas auxin promotes cell divisions in the cambium. Loss of function mutations in cambium-related genes similarly affected tissue attachment and vascular reconnection. Together, our findings highlight the importance and function of cambium while providing evidence to refine our understanding of graft development.

## Results

### Establishing a tissue-specific inducible *bdl* system

Auxin response is important for successful graft formation (Melnyk et al., 2015) and we confirmed this by treatment with the auxin receptor inhibitor auxinole and by grafting with the *bdl-2* mutant, both of which reduced phloem and xylem reconnection at the graft junction (Figure 1A-D). While *bdl* had a strong suppressive effect on grafting, the native BDL promoter drives *bdl* expression broadly in the stele (Hamann et al., 2002) suggesting any one of these cell types might be relevant for graft formation. To better understand where auxin signaling is required, we employed a previously published two-component inducible tissue-specific expression system (Schürholz et al., 2018). We cloned the *bdl-2* gene and drove it under a synthetic promoter (“effector line”) that activates only in the presence of a cognate transcription factor and crossed the resulting transgenic plant into 11 previously published lines that express the transcription factor in an inducible and tissue specific manner in various root cell types (“driver line”)(Figure 1E, Figure S1A). Upon dexamethasone (DEX) induction of the resulting F1 plants, we observed tissue-specific *mTurquoise2* fluorescence (Figure 1F) consistent with cell-specificity of the transgene. DEX treatment of F1 plants increased hypocotyl lengths in plants expressing *bdl* in the procambium (*pATHB8* and *pPXY*) and xylem pole pericycle (*pXPP*), while it decreased primary root growth when *bdl* was expressed in the epidermis (*pML1*)(Figure 2A-C). Several lines showed an increase in lateral root numbers when *bdl* was misexpressed including in the endodermis (*pSCR*), phloem precursors (*pSMXL5*), procambium (*pPXY, pWOX4*), xylem pole pericycle (*pXPP*) and xylem precursors (*pTMO5*)(Figure 2D). Such effects were consistent with perturbed auxin signaling similar to what has been observed in other auxin signaling mutants (Lincoln et al., 1990; Leyser, 2017).

**Figure 1.**
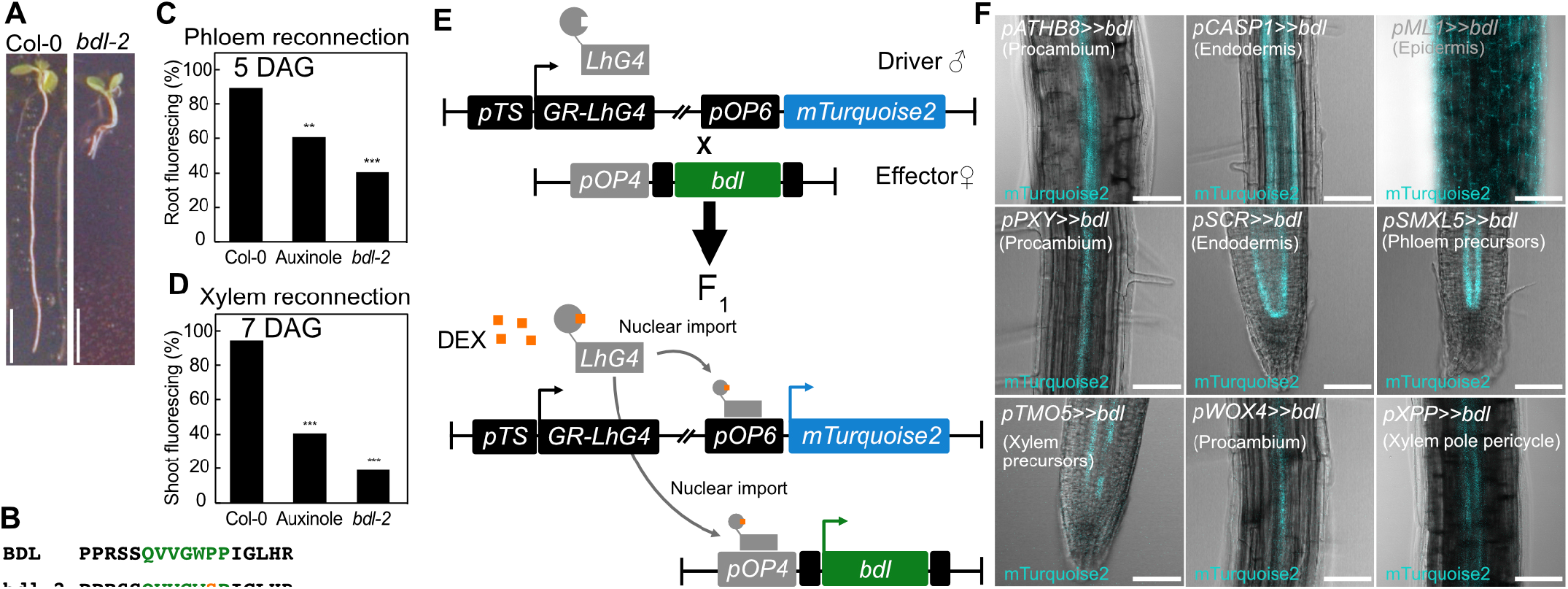
The *bdl* mutation is relevant for graft formation and for using in a cell-specific misexpression system. (A-B) *bdl* auxin resistance phenotype is caused by a point mutation resulting in a praline 74 to serine amino acid change in the conserved degradation domain. Scale bar is 1 cm. (C-D) Proportion of grafted *Arabidopsis* that transported CFDA to the rootstock at 5 days after grafting (C) or to the scion at 7 days after grafting (D). 30 uM auxinole was used. n=20 per treatment. ^*^p<0.05; ^**^p<0.01; ^***^p<0.001; Fisher’s exact test compared to mock controls. (E) The design of inducible cell-type specific expression system. The *bdl* effector line was crossed with driver lines expressing a synthetic transcription factor GR-LhG under various tissue-specific promoters *(pTS)*. In the resulted F_1_ seedlings, GR-LhG drives the expression of *bdl* as well as the *mTurquise2* reporter when induced by DEX. (F) Validation of tissue specific expression of *bdl* in the targeted tissues by confocal imaging and detection of mTurquise2 signals. Scale bar is 100 μm.

**Figure 2.**
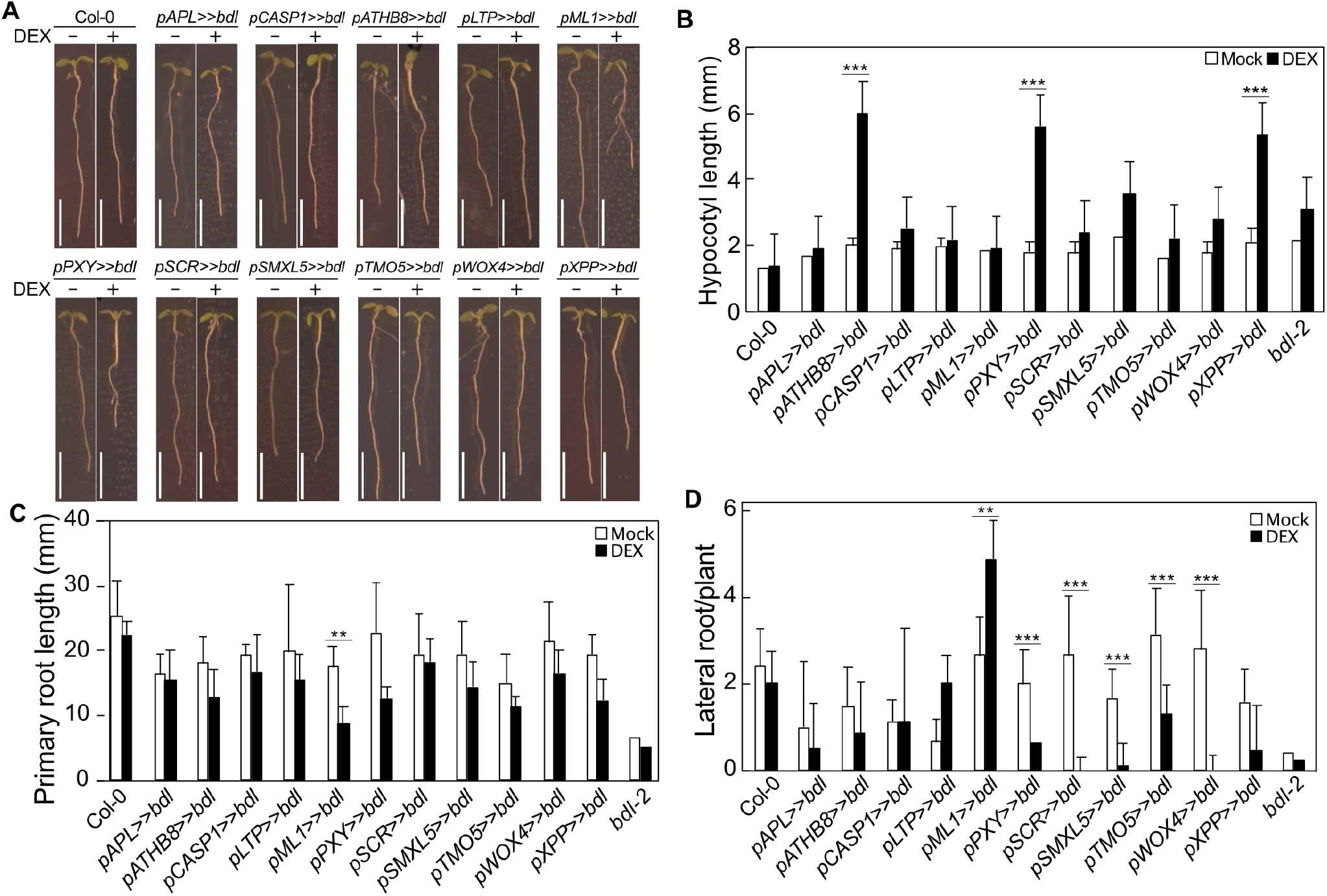
Perturbing cell-specific auxin responses modifies hypocotyl and root phenotypes. (A-D) Phenotypic analysis of the F_**1**_ plants expressing *bdl* in various tissues upon DEX induction. Measurement of hypocotyl length (B), primary root length (C), and lateral root number (D) of transgenic plants expressing *bdl* under various tissue-specific promoter. Values represent mean±s.d. n= 20-24 plants per genotype per treatment. ^*^p<0.05; ^**^p<0.01; ^***^p<0.001; student’s t-test compared to mock controls. Scale bar is 1 cm.

### Auxin response in the procambium is required for successful graft formation

Successful grafting involves tissue attachment followed by callus formation, phloem reconnection and xylem reconnection (Melnyk et al., 2015). To understand the role of individual cell types in these processes, we performed attachment and phloem reconnection assays on the misexpression lines. Only those plants expressing *bdl* in the procambium (*pATHB8* and *pPXY*) failed to attach, form callus, reconnect phloem and reconnect xylem (Figure 3A-B, S1C-D). However, *bdl* misexpression in the xylem pole pericycle cells (*pXPP*) and phloem precursors (*pSMXL5*) specifically inhibited phloem reconnection (Figure 3A-B, Figure S1). To exclude the possibility that this effect was specific to *bdl*, we also misexpressed *iaa18* and found similar effects on phloem reconnection with procambium and phloem precursor lines (Figure S1B). Given the strong effects of cell specific expression of *bdl* upon grafting, we tested the importance of *bdl* misexpression in the scion compared to the rootstock. While *pATHB8* misexpression in the rootstock and scion reduced phloem reconnection, *pPXY* misexpression affected phloem reconnection only when in the scion (Figure 3C,E). Inhibition of tissue attachment occurred with *pATHB* misexpression in the rootstock, while *pATHB* and *pPXY* misexpression in both rootstock and scion was needed for the strongest block in tissue attachment (Figure 3C,E). Thus, although tissue adhesion was needed for vascular reconnections, by using heterografting we could uncouple these two processes to show a role for procambium in both attachment and phloem reconnection. To further test the requirements of cambium, we grafted plants and allowed them to attach and develop callus before inducing *bdl* for various days in the procambium. Blocking auxin responses in procambium from 0 to 1 day after grafting (DAG) blocked attachment. Blocking auxin response 1-2 DAG allowed tissue attachment but inhibited phloem reconnection (Figure 3D,F). Induction at 3 DAG had no effect on phloem reconnection since phloem was formed by then (Figure 3D,F). Together, our data revealed a role for auxin signaling in the procambium driving both tissue attachment and phloem formation.

**Figure 3.**
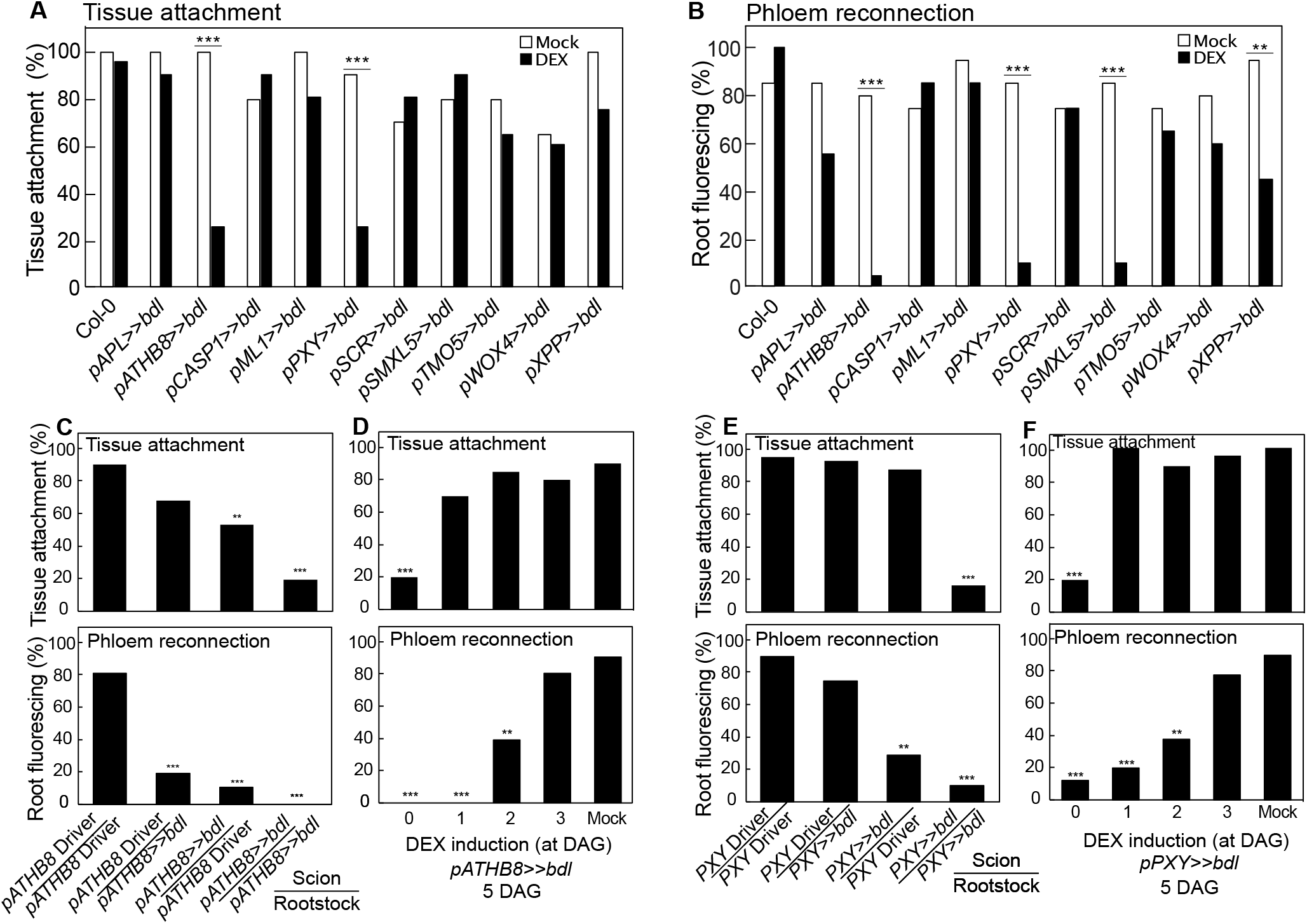
Perturbing auxin responses in the procambium inhibited successful graft formation. (A) Proportion of grafts attached in transgenic *Arabidopsis* misexpressing *bdl* at 5 days after grafting (DAG). (B) Proportion of grafted transgenic *Arabidopsis* that transported CFDA to rootstocks at 5 DAG. (C,E) Proportion of DEX-treated and grafted *pATHB8>>bdl* (C) or *pPXY>>bdl* (E) to the respective driver lines that attached or transported CFDA. The homo- and hetero-graft combinations are indicated. All plants were sampled at 5 days after grafting. (D,F) Proportion of grafts attached or transported CFDA to the rootstock of grafted *pATHB8>>bdl* (D) or *pPXY>>bdl* (F). DEX was applied at indicated time points. Plants were sampled at 5 days after grafting. (A-F) n=20-30 per treatment. ^*^p<0.05; ^**^p<0.01; ^***^p<0.001; Fisher’s exact test compared to mock or driver controls.

### Auxin responses in procambial tissues are required for cell expansion and cell division

To understand how *bdl* blocked graft formation, we examined graft junctions misexpressing *mTurquoise2* with *pATHB8, pSMXL5* and *pPXY* in the presence or absence of *bdl*. In driver lines, we observed the domain of *pATHB8* and *pSMXL5* expression in the vasculature and the callus tissues connecting the scion and rootstock (Figure 4A,C). *pPXY* expression was more diffuse and appeared in the vasculature but also outer cell layers around the graft junction including the endodermis and cortex layers (Figure 4B). In plants expressing both driver and *bdl* effector, *mTurquoise2* signal was noticeably reduced with little signal present at the cut site and the vascular diameters reduced (Figure 4D-G). *pSMXL5* misexpressing *bdl* plants remained attached though this appeared through non *SMXL5* expressing cells, likely instead cortex or epidermal tissues (Figure 4F). We also imaged cut but non-grafted hypocotyls to better observed the effects of *bdl* misexpression at the cut. Cut top expression patterns for *pATHB8, pSMXL5* and *pPXY* driver lines were similar as those in the scion, but cut root expression patterns showed little cell division and cell expansion compared to the grafted rootstock (Figure S2A-C, S2G-I). Thus, it appeared that contact between rootstock and scion was necessary for vascular proliferation. In plants expressing both driver and *bdl* effector, the *mTurquoise2* expression domain was greatly reduced and limited to the vascular tissues which showed little proliferation and expansion compared to the driver lines (Figure S3D-I), consistent with *bdl* misexpression blocking vascular proliferation, cell expansion and callus formation in the grafted or cut plants. We further analysed the role of cell division by staining with the fluorescent labeled thymidine-analogue EdU that incorporates into newly synthesized DNA of dividing cells (S phase). We detected high EdU staininig in grafted driver tissues, particularly in the vasculature, consistent with high levels of active cell division in these tissues (Figure 4H). Conversely, EdU signals were undetectable in the graft junction of plants expressing *bdl* in procambium and phloem precursors (Figure 4H), implying that *bdl* suppressed cell division. Moreover, grafted plants expressing *bdl* in phloem precursors attached their tissues but with little cell division and no vascular connection (Figure 4H). Our data suggest that graft failures induced by *bdl* expression were due in part to the inhibition of procambial cell division that prevented callus and phloem formation.

**Figure 4.**
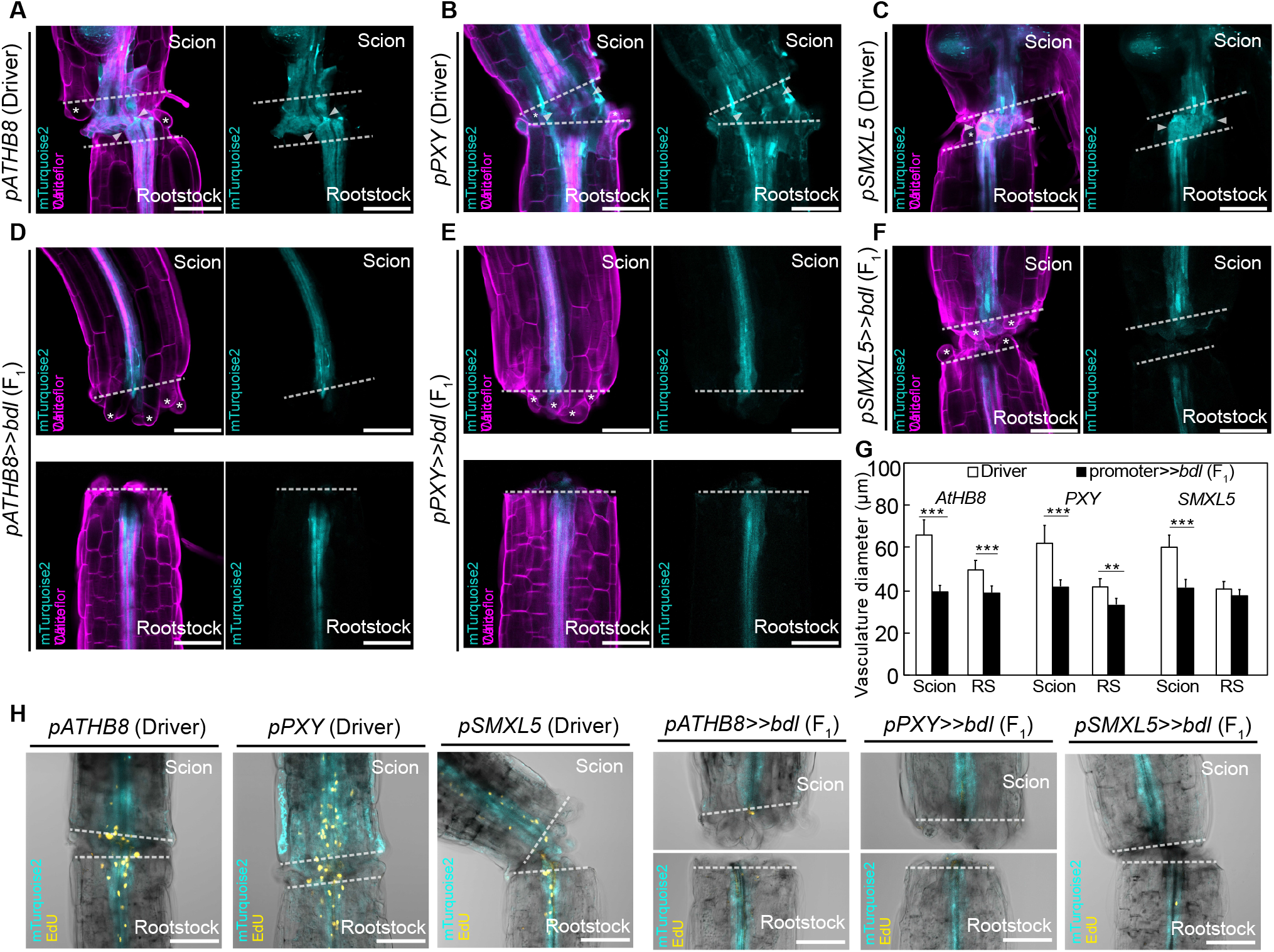
Auxin response is important for cambial expansion and cell divisions. Longitudinal optical sections of DEX-treated grafted hypocotyl driver lines (A-C) and *bdl* missexpression (D-F) at 5 days after grafting (DAG). The genotypes are indicated. White asterisks indicate expanding cortex cells. White arrow heads indicate vascular connection in the graft junction. AR: adventitious root. (G) Vasculature diameter including pericycle, cambium, xylem, and phloem of grafted plants of indicated genotype at 5 DAG, 100 μm from the cut surface. (mean±s.d.; n=11-16 plants per genotype. ^*^p<0.05; ^**^p<0.01; ^***^p<0.001; student’s t-test compared to driver lines). (H) EdU staining detecting cell proliferation at graft junctions of plants expressing *bdl* in various cell types. The genotypes are indicated. Dashed lines indicate the cut site; bars=100 μm.

Our *bdl* misexpression analysis suggested an important role for the procambium and phloem precursors in graft attachment and phloem reconnection. To further investigate these roles, we grafted mutants in these cell types including cambium (*wox4* and *pxy,wox4*), phloem precursors (*smxl4,5*), xylem pole pericycle (*slr1*) or endodermis/cortex (*shr*). Cambium-related mutants reduced attachment and phloem reconnection, whereas the *slr1* and *smxl4,5* mutants showed a reduction in phloem reconnection. *shr* however did not have significant changes in attachment or reconnection (Figure 5A-D). Together, it appeared that these loss of function mutants were consistent with our *bdl* misexpression analysis revealing a critical role for the cambium and procambium.

**Figure 5.**
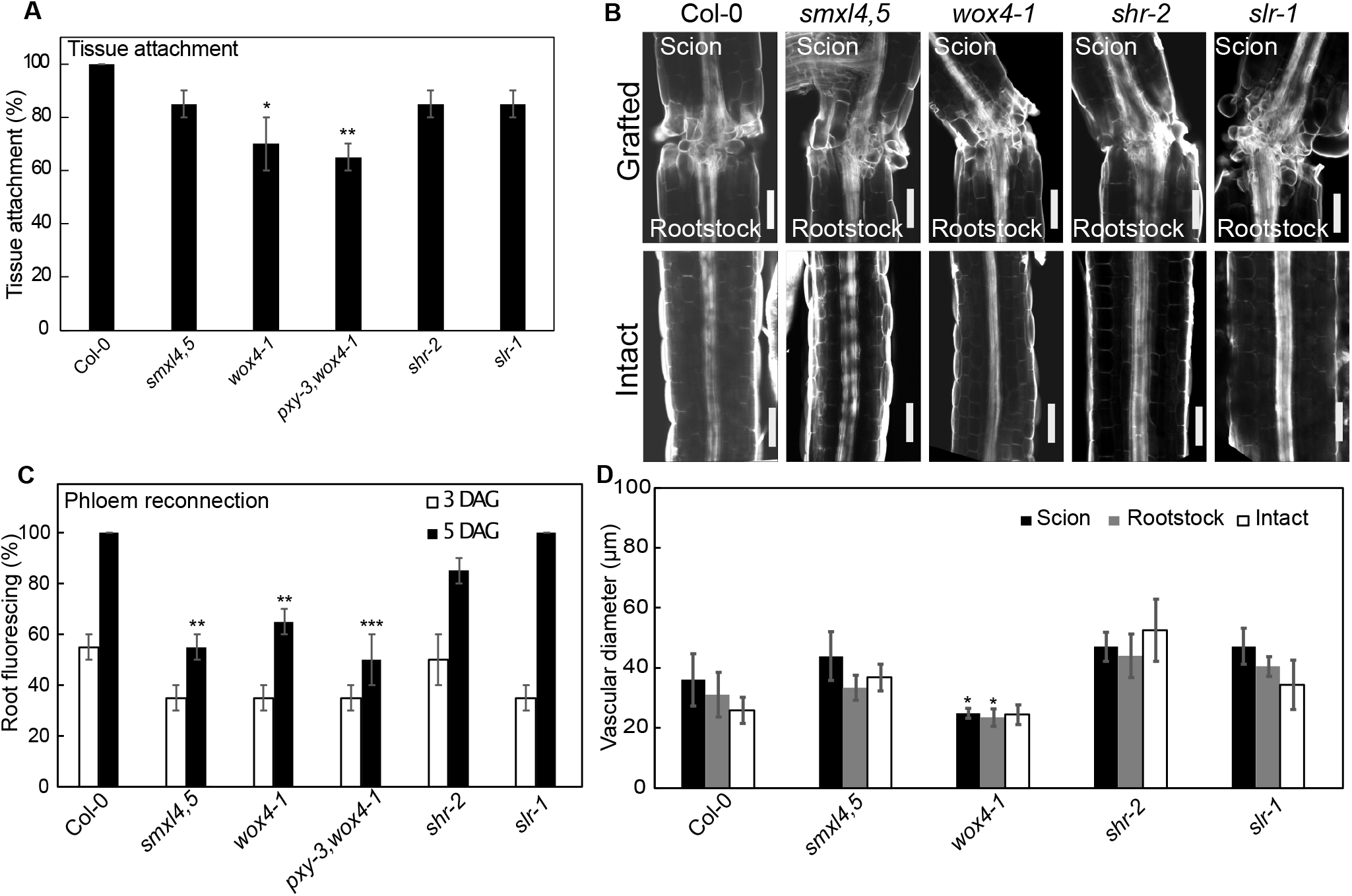
Cambial mutants are impaired in tissue attachment and phloem reconnection. (A) Tissue attachment in *Arabidopsis* mutants and wildtype 1.5 days after grafting. The mean + SEM from 20 plants per genotype is shown. (Fisher’s exact test; ^*^p<0.05, ^**^p<0.01) (B) Longitudinal optical sections of grafted or intact cambial mutants five days after grafting. Scale bar is 100 μm. (C) Phloem reconnection in *Arabidopsis* mutants and wildtype 3 and 5 days after grafting. The mean + SEM from 17-20 plants per genotype is shown. (Fisher’s exact test; ^**^p<0.01, ^***^p<0.001) (D) Vasculature diameters of hypocotyls in *Arabidopsis* mutants and wildtype 5 days after grafting. The measurement includes pericycle, cambium, xylem and phloem of grafted scion and root stock and of intact plants. The mean + SD from 5-19 plants per genotype is shown. Student’s t-test compared to Col-0; ^*^p<0.05.

## Discussion

Successful graft formation employs multiple cellular regeneration processes including tissue adhesion, callus formation and vascular reconnection. Here, we used an inducible cell-type specific expression system to investigate the role of individual cell types during graft formation in a spatio-temporal and non-destructive manner. We consistently observed that perturbing auxin signaling in procambium during graft formation inhibited cell division, graft attachment, phloem differentiation and xylem differentiation. Procambium appears to have separate roles for attachment and vascular differentiation since, by inducibly activating *bdl*, we could allow attachment to occur yet block phloem and xylem reconnection. Thus, our data suggest auxin signaling in procambium was critical for attachment and phloem reconnection, and phloem formation likely occurred via a *AtHB8* and *PXY* expressing domain in the graft junction. Notably, *pSMXL5>>bdl* plants could reconnect xylem (Fig S1) despite their inability to reconnect phloem (Fig 3B), suggesting that phloem reconnection is not required for xylem reconnection, and auxin signaling in the *SMXL5* domain was not necessary for tissue attachment and callus formation. We observed some spreading of these domains outside of the cambial area, particularly for *pPXY*, and our analysis is also limited to where the promoter is active rather than the origin of the cells. However, our genetic analysis showed a reduction in attachment and phloem transport rates for cambial mutants, consistent with our *bdl* analysis and consistent with a previous study reporting the requirement of auxin signaling for procambium differentiating into phloem precursors during plant radial growth (Smetana et al., 2019). *PXY* gene expression is also required for procambial cells to maintain the polarity of cell division by programing the cells into the phloem lineage during vascular development (Fisher & Turner, 2007; Smit et al., 2019). Notably, our analysis revealed that procambium (*pATHB8*), procambium on the phloem side (*pSMXL5*) and procambium on the xylem side (*pPXY*) impaired grafting when auxin signaling was blocked yet other markers expressed in the procambium (*pWOX4*) or developing xylem (*pTMO5*) showed no effect. It’s possible that these domains are not directly affected by a block in auxin signaling or due to functional redundancy show weaker effects.

Our analyses revealed auxin responsive cell types but also shed light on the importance of rootstock or scion specific response. While *pATHB8* driven *bdl* expression affected both rootstocks and scions, *pPXY* had its strongest effects in the scion. Previous studies have shown that mutating auxin related genes such as *ALF4, AXR1* and *BDL* all have their strongest effects in the rootstock, while auxin biosynthesis genes such as YUCCAs are required in the scion (Melnyk et al., 2015; Serivichyaswat et al., 2022). These contrasting findings suggest that where auxin is perceived in the rootstock or scion is critical to whether grafts succeed or fail. A broadly expressed stele promoter such as *pATHB8* might thus have an ability to perturb a greater degree of auxin response. In support of the notion that both rootstock and scion auxin response are relevant, mutating DOFs necessary for cambial divisions in the rootstock or scion both perturbs grafting (Zhang et al., 2022). A role for cambium in grafting has been long speculated including the advice to align cambiums for grafting success in horticulture (Garner & Bradley, 2013). Tomato *WOX4* mutants and Arabidopsis DOF transcription factor mutants important for cambial divisions both perturb graft formation (Thomas et al., 2021; Zhang et al., 2022). Thus, our findings contribute to deepening our understanding of the role of cambium and how this tissue is critical for grafting success.

## Materials and Methods

### Cloning

Plasmid construction was done using modules of Greengate cloning (Lampropoulos et al., 2013). For the entry module for *bdl*, the CDS of *bdl* was amplified from cDNA of the *bdl-2* mutant, then inserted into pGGC000 to create pGGC-bdl. The restriction and ligation reactions were done using BsaI-HF and T4 ligase (NEB), respectively. The resulting plasmids were transformed into *E. coli* using chemically competent cells (Subcloning EfficiencyTM DH5α Competent Cells, ThermoFisher Scientific) according to the manufacturer’s protocol. The transformed cells were cultured and selected on LB medium with 100 μg/mL ampicillin. The plasmids were extracted using Plasmid DNA Miniprep Kit (ThermoFisher), and the sequences were confirmed by sequencing of the ligation sites (Macrogen). The final plant transformation vector pTRPY-*bdl* was created by Greengate reaction as previously described (Lampropoulos et al., 2013), using pGGA016 (pOP plant promoter), pGGB003 (B-dummy), pGGC-bdl (bdl CDS), pGGD002 (D-dummy), pGGE009 (UBQ10 terminator), pGGF008 (pNOS::Basta resistance::tNOS), and pGGZ003 (empty destination vector). The ligation product was used for *E. coli* transformation, then the cells were cultured on LB medium with 100 μg/mL spectinomycin. The sequence was initially confirmed by digestion analysis, then sequencing of the ligation site. The plasmid was then co-transformed with helper plasmid pSOUP in electrocompetent *Agrobacterium tumefaciens* strain GV3101 then the cells were cultured on LB medium with 100 μg/mL spectinomycin.

The *bdl* effector line was generated by introducing pTRPY-*bdl* into *Arabidopsis thaliana* Col-0 using the floral dip method (Clough & Bent, 1998). The T_3_ homozygous transgenic plants were selected on ½MS media, 1% plant agar, pH 5.8, containing 7.5 mg/mL Basta™ (glufosinate ammonium) and used for the subsequent experiments.

### Plant materials, growth conditions, and grafting

All *Arabidopsis thaliana* (L.) used throughout this study was in Columbia (Col-0) background unless otherwise indicated. Seeds of effector lines (Schürholz et al., 2018) purchased from NASC included *pAPL* (N2107943), *pATHB8* (N2107955), *pCASP1* (N2107953), *pML1* (N2107954), *pPXY* (N2107941), *pSCR* (N2107940), pSMXL5 (N2107942), *pTMO5* (N2107952), *pWOX4* (N2107945), and *pXPP* (N2107938). Mutant lines used included *bdl-2* (Hayward et al., 2009), *slr-1, shr-2, wox4-1* and *smxl4,5*. The F_1_ seeds were obtained by crossing between the effector lines and the *bdl* effector line. For *in vitro* germination, seeds were surface sterilized with 70% (v/v) ethanol for 10 minutes, followed by 90% (v/v) ethanol for 10 minutes. The seeds were then sown and germinated on ½MS media, 1% plant agar, pH 5.8. After stratification in the dark at 4°C overnight, the seeds were transferred to 20°C short-day growth conditions (8 h of 140 μmol m^-2^ s^-1^). For DEX induction treatment, F_1_ seeds were sown and germinated without DEX, and five-day-old seedlings were transferred to media containing 5μM DEX (Sigma-Aldrich) for five days before phenotypic analyses. *Arabidopsis* grafting was performed on five-day-old seedlings using previously published protocols (Melnyk, 2017a; Melnyk, 2017b). For DEX induction, 5μM DEX or equal volume of DMSO (Fisher Scientific) was used instead of water in the graft setting. Plants were sampled for phloem and xylem reconnection assays at 5 and 7 days after grafting, respectively.

### Phloem and xylem connection assays

Phloem and xylem transport rates were measured using a previously published protocol (Serivichyaswat et al., 2022).

### Cell proliferation detection

EdU staining was performed using Click-iT™ EdU Cell Proliferation Kit (Invitrogen). In brief, 10μM EdU were applied to grafted plants (3 DAG) for 1 hr followed by tissue fixation, permeabilization, staining following the manufacturer’s instruction. Visualization of EdU was performed with confocal microscopy (Zeiss LSM780).

### Confocal imaging

Graft junctions were imaged using Zeiss LSM780 NLO laser scanning confocal microscope. The tissues were cleared and stained with Calcoflour White staining protocol (Ursache et al., 2018), and visualised on the LSM780 with 405 nm excitation, 5% laser power, 410-529 nm emission, and 210 PMT. The mTurquoise2 signals were detected with 458 nm excitation, 5% laser power,460-516 nm emission. Vascular diameter quantifications included cambium, xylem, phloem and pericycle tissues, and measured the distance between the pericycle layers encompassing the vascular bundle 100-200 μm above the cut site. The images were processed and analyzed using FIJI software.

### Accession Numbers

The Arabidopsis Genome Initiative numbers of genes used in this study are as follows: *SCR* (AT3G54220), *ATHB8* (AT4G32880), *XPP* (At4g30450), *PXY* (AT5G61480), *TMO5* (AT3G25710), *SMXL5* (AT5G57130), *CASP1* (AT2G36100), *APL* (AT1G79430), *WOX4* (AT1G46480), *ML1* (AT4G21750), *SHR* (AT4G37650), *SLR* (AT4G14550), *SMXL4* (AT4G29920), *SMXL5* (AT5G57130), *WOX4* (AT1G46480) and *BDL* (AT1G04550).

## Acknowledgements

We thank the Nottingham Arabidopsis Seed Center (NASC), Addgene, Jan Lohmann (University of Heidelberg, Germany) and Thomas Greb (University of Heidelberg, Germany) BRC for sharing materials. P.T.S. and C.W.M were supported by a Wallenberg Academy Fellowship (2016-0274). M.F. and C.W.M were supported by a European Research Council starting grant (GRASP-805094). A.K. was supported by a MSCA Postdoctoral Fellowship (UMOCELF - 101069157).

## Figure legends

**Supplemental figure 1.** (A) *bdl* coding sequence was cloned into the Greengate cloning system. Six entry plasmids, including a synthetic *pOP* promoter, an empty N-terminal tag, a *bdl*-coding sequence, an empty C-terminal tag, a UBQ10-terminator, and a plant selectable marker Basta resistance cassette, were used for a Greengate reaction, yielding the plant transformation destination vector pTRPY_*pOP4::bdl*, which was subsequently used to generate a transgenic *bdl* effector line. (B) Proportion of grafted *Arabidopsis* misexpressing *iaa18* that transported CFDA to the rootstock at 5 days after grafting. (C) Hypocotyl callus size from selected genotypes 5 days after cutting. mean±s.d., n= 20-22 hypocotyl per genotype and treatment. ^*^p<0.05; ^**^p<0.01; ^***^p<0.001; student’s t-test compared to mock controls. (D) Callus formation from hypocotyls of cut but ungrafted plants expressing bdl in procambium (*pATHB8* and *pPXY*). Scale bars=500 μm. Dashed lines indicate the cut site. (E) Proportion of grafted transgenic *Arabidopsis* of selected genotypes that transported CFDA to scions. n=20-25 plants per genotype per treatment. ^*^p<0.05; ^**^p<0.01; ^***^p<0.001; Fisher’s exact test compared to mock controls. (F) Proportion of grafted *pATHB8>>bdl* or *pPXY>>bdl* that transported CFDA to the scion. DEX was applied at 3 days after grafting and plants were sampled at 7days after grafting. n=20-30 per treatment. ^*^p<0.05; ^**^p<0.01; ^***^p<0.001; Fisher’s exact test compared to mock controls.

**Supplemental figure 2. Auxin responses in cambium is required for tissue expansion at the cut site**. Longitudinal optical sections of DEX-treated and cut but ungrafted hypocotyl driver lines (A-C) and *bdl* misexpression lines (D-F). The genotypes are indicated. White asterisks indicate expanding cortex cells. AR: adventitious root. Dashed lines indicate the cut site; bars=100 μm. (G-I) Vasculature diameter including pericycle, cambium, xylem, and phloem of cut but ungrafted plants of indicated genotype, 100 μm from the cut surface (mean±s.d.; n=15-18 plants per genotype. ^*^p<0.05; ^**^p<0.01; ^***^p<0.001; student’s t-test compared to driver lines). All plants were sampled at 5 days after cutting.

