## Supplemental Figures 1-2 for "Auxin signaling in the cambium promotes tissue attachment and vascular development during *Arabidopsis thaliana* graft formation"

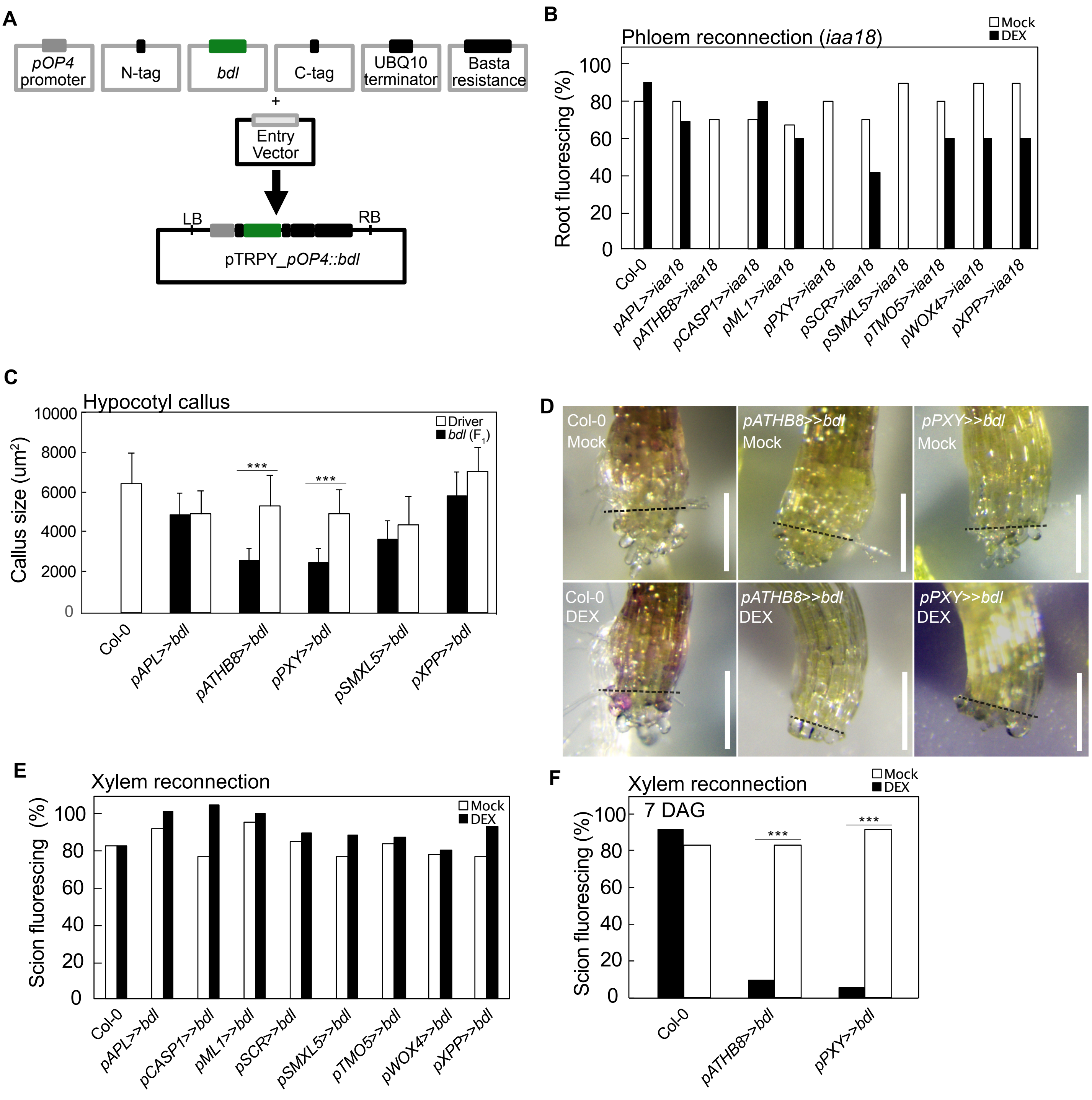


**Supplemental figure 1.** (A) *bdl* coding sequence was cloned into the Greengate cloning system. Six entry plasmids, including a synthetic *pOP* promoter, an empty N-terminal tag, a *bdl*-coding sequence, an empty C-terminal tag, a UBQ10-terminator, and a plant selectable marker Basta resistance cassette, were used for a Greengate reaction, yielding the plant transformation destination vector pTRPY_*pOP4::bdl*, which was subsequently used to generate a transgenic *bdl* effector line. (B) Proportion of grafted *Arabidopsis* misexpressing *iaa18* that transported CFDA to the rootstock at 5 days after grafting. (C) Hypocotyl callus size from selected genotypes 5 days after cutting. mean±s.d., n= 20-22 hypocotyl per genotype and treatment. *p<0.05; **p<0.01; ***p<0.001; student's t-test compared to mock controls. (D) Callus formation from hypocotyls of cut but ungrafted plants expressing bdl in procambium (*pATHB8* and *pPXY*). Scale bars=500 μm. Dashed lines indicate the cut site. (E) Proportion of grafted transgenic *Arabidopsis* of selected genotypes that transported CFDA to scions. n=20-25 plants per genotype per treatment. *p<0.05; **p<0.01; ***p<0.001; Fisher's exact test compared to mock controls. (F) Proportion of grafted *pATHB8>>bdl* or *pPXY>>bdl* that transported CFDA to the scion. DEX was applied at 3 days after grafting and plants were sampled at 7days after grafting. n=20-30 per treatment. *p<0.05; **p<0.01; ***p<0.001; Fisher's exact test compared to mock controls.

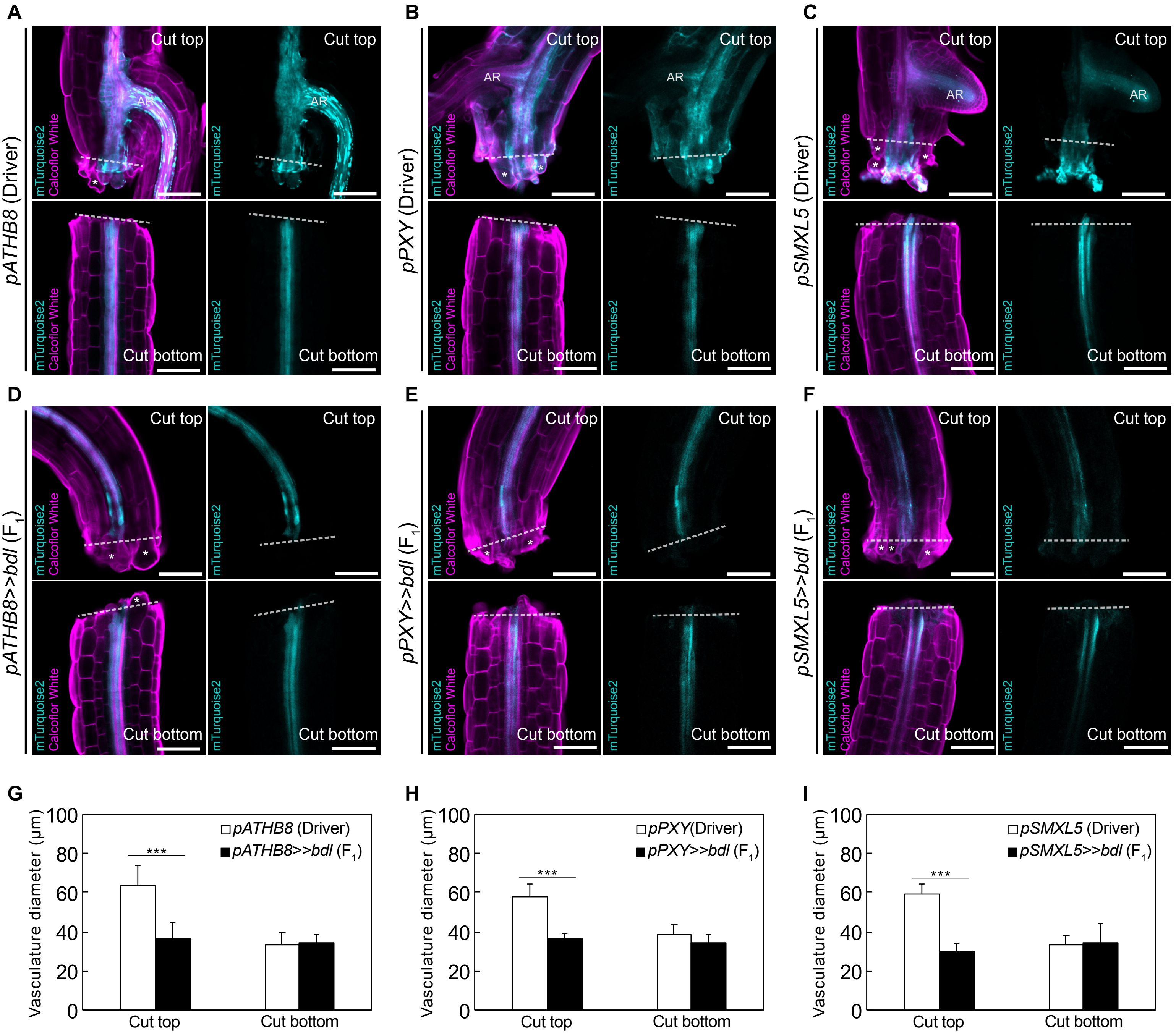


**Supplemental figure 2. Auxin responses in cambium is required for tissue expansion at the cut site.** Longitudinal optical sections of DEX-treated and cut but ungrafted hypocotyl driver lines (A-C) and *bdl* misexpression lines (D-F). The genotypes are indicated. White asterisks indicate expanding cortex cells. AR: adventitious root. Dashed lines indicate the cut site; bars=100 μm. (G-I) Vasculature diameter including pericycle, cambium, xylem, and phloem of cut but ungrafted plants of indicated genotype, 100 μm from the cut surface (mean±s.d.; n=15-18 plants per genotype. *p<0.05; **p<0.01; ***p<0.001; student's t-test compared to driver lines). All plants were sampled at 5 days after cutting.
